# Glucagon-like peptide-1 receptors in the gustatory cortex influence food intake

**DOI:** 10.1101/2022.08.26.505475

**Authors:** Amanda M. Dossat, Milayna Kokoska, Jessica Whitaker-Fornek, Aishwarya S. Kulkarni, Erica S. Levitt, Daniel W. Wesson

## Abstract

The gustatory region of the insular cortex (GC) processes taste information in manners important for taste-guided behaviors, including food intake itself. In addition to oral gustatory stimuli, GC activity is also influenced by physiological states including hunger. The specific cell-types and molecular mechanisms that afford with GC with such influences on food intake are unclear. Glucagon-like peptide 1 (GLP-1) is produced by neurons in the brain whereafter it can act upon GLP-1 receptor-expressing (GLP-1R+) neurons found in several brain regions. In these brain regions, GLP-1R agonism suppresses homeostatic food intake and dampens the hedonic value of food. Here, we report in mice of both sexes that cells within the GC express GLP-1R mRNA and further, by *ex vivo* brain slice recordings, that GC GLP-1R+ neurons are depolarized by the selective GLP-1R agonist, exendin-4 (Ex-4). Next we found that chemogenetic stimulation of GLP-1R+ neurons, and also pharmacological stimulation of GC-GLP-1Rs themselves, both reduced homeostatic food intake. When maintained on a high-fat diet, obese mice exhibited impaired food intake responses when Ex-4 was administered into the GC. Yet, when obese mice were switched to a low-fat diet, the effect of GC Ex-4 was restored – indicating that GC GLP-1R influences may depend upon palatability of the food. Together, these results provide evidence for a specific cell population in the GC which may hold roles in both homeostatic and hedonic food intake.

## Introduction

The gustatory region of the insular cortex (GC) is a brain region wherein tastant information is processed in manners critical for taste-guided behaviors, including food intake and the development of taste preferences. In line with this role, GC activity in animals ranging from mice, rats, to humans, can reflect the perceived palatability of tastes (Katz et al., 2001; Small et al., 2001; Accolla and Carleton, 2008; Sadacca et al., 2012, 2016; Jezzini et al., 2013; Bouaichi and Vincis, 2020; Vincis et al., 2020). The GC is a highly innervated cortical region (Gehrlach et al., 2020), that receives projections from gustatory and visceral-associated brain regions (Livneh et al., 2017, 2020; Di Lorenzo, 2021). The GC also receives input from brain areas that may influence the desire to eat (Di Lorenzo, 2021) and importantly projects to brain areas involved in regulating food intake (Yasui et al., 1991; Zhang et al., 2009; Wu et al., 2020; Samuelsen and Vincis, 2021).

In addition to orosensory information, the GC is also sensitive to metabolic status, wherein GC neurons are more active during hunger *vs*. satiety (Small et al., 2001; Hinton et al., 2004; de Araujo et al., 2006; Malik et al., 2008; Livneh et al., 2017). Beyond processing taste and interoceptive information, manipulation of GC activity alters ingestive behavior (Baldo et al., 2016; Stern et al., 2020, 2021; Wu et al., 2020; Zhang-Molina et al., 2020). Taken together, the GC is positioned to modulate ingestive behavior *via* integration of gustatory and visceral signals (Katz et al., 2001, 2002; Maffei et al., 2012). While there is both preclinical and clinical evidence supporting the role of the GC as a key interoceptive brain region, the cellular mechanisms remain poorly understood.

Taste-related information is transmitted to the brain *via* signals arising from the mouth, while meal-related interoceptive information is conveyed *via* signals from within the gut. This concert of signals allows one to experience hunger/fullness and control ingestion. There are many peptides which influence ingestive behavior (Crespo et al., 2014). For example, the gut signals fullness (*e*.*g*., gastric stretch) through the secretion of meal-related peptides such as GLP-1 which binds to GLP-1Rs on the vagus nerve (Williams et al., 2016). The nucleus of the solitary tract (NTS) contains a population of GLP-1-producing neurons that are activated by gastric stretch (Vrang et al., 2003), leptin (Hisadome et al., 2010), and cholecystokinin (Hisadome et al., 2011). These neurons send projections to feeding-relevant brain regions that express GLP-1 receptors (GLP-1R) (Larsen et al., 1997; Merchenthaler et al., 1999; Vrang et al., 2007), which include the hypothalamus (Larsen et al., 1997; Rinaman, 2010), nucleus accumbens (Dossat et al., 2011; Alhadeff et al., 2012), and the lateral septum (Terrill et al., 2019), among others, wherein GLP-1R agonists reduce food intake (e.g., (Tang-Christensen et al., 1996; McMahon and Wellman, 1998; Dossat et al., 2011; Kanoski et al., 2011; Alhadeff et al., 2012). The precise mechanism through which GLP-1R agonism influences ingestive behavior seems to depend in-part upon the brain region wherein GLP-1Rs are expressed. For example, hypothalamic GLP-1R activation suppresses homeostatic food intake (McMahon and Wellman, 1998), while GLP-1R activation within the nucleus accumbens dampens the hedonic value of food (Dickson et al., 2012; Dossat et al., 2013; López-Ferreras et al., 2018). The food intake-reducing effects of GLP-1R agonism is influenced by consumption and maintenance on a high-fat diet (Williams et al., 2011; Duca et al., 2013). During investigations into other brain regions, we serendipitously observed expression of *Glp1r* mRNA in the GC, which raised the exciting possibility that GLP-1 acts upon GC neurons to afford its well-established influence on ingestive behavior.

Here we used a combination of fluorescent *in situ* hybridization, *ex vivo* electrophysiology, chemogenetics, behavioral pharmacology, and diet-induced obesity mouse models to investigate a possible role for GLP-1Rs in the GC. We aimed to answer the following questions: Where are GLP-1R expressed in the GC, are there anatomical or cortical layer-specific expression of GLP-1Rs? Are GLP-1R+ neurons responsive to GLP-1R ligands? Does activation and/or inhibition of GC GLP-1Rs influence food intake? And finally, due to the known application of GLP-1R analogues as therapeutics for obesity (e.g., (Wilding et al., 2021)), does the possible influence of these receptors change in the context of obesity?

## Materials and Methods

### Animals

Adult C57BL/6J mice (N=51; 27 males, 18 females) (bred in a University of Florida vivarium from breeder stock originating from The Jackson Laboratory; Bar Harbor, ME), Glp1r^tm1.1(cre)Lbrl^ heterozygotes (Williams et al., 2016) (N=14; 8 males, 6 females; “Glp1r-Cre”; The Jackson Laboratory, strain #029283), and Glp1r-Cre heterozygotes crossed with homozygotic B6.Cg-Gt(ROSA)26Sortm9(CAG-tdTomato)Hze/J mice (Madisen et al., 2010) (N=17; 19 males, 6 females, “Glp1r-Cre;Ai9”; The Jackson Laboratory, strain #007909) were used in the present study. Mice were housed in a temperature-controlled vivarium on a 12:12h light:dark cycle with *ad libitum* access to water and rodent chow (catalog #2018, 2918, Teklad, Envigo, Indianapolis, IN) except where otherwise stated. All of the experimental procedures were conducted in accordance with the guidelines from the National Institutes of Health and approved by th University of Florida Institutional Animal Care and Use Committee.

### RNAscope® Fluorescent in situ hybridization (FISH)

C57BL/J mice mice were anesthetized with Fatal Plus (100 mg/kg, catalog #00298-9373-68, Vortech Pharmaceuticals, Dearbor, MI) and transcardially perfused with chilled 0.9% saline. Brains were quickly removed (<5 min), flash-frozen in isopentane (catalog #AC126470010, Fisher Scientific, Waltham, MA) on dry ice, and stored at -80°C until further processing. Coronal sections (10µM) were cryosectioned at -20°C, mounted onto Super Frost Plus slides (catalog #12-550-15, Fisher Scientific), and FISH experiments were performed according to the manufacturer’s protocol for fresh-frozen samples using the RNAscope Multiplex Fluorescent v2 Assay (catalog #323136, Advanced Cell Diagnostics, Newark, CA) (Wang et al., 2012). The probe for *Glp1r* (catalog #418851) was paired with the Opal Fluorophore 690 (catalog #NC1605064, Akoya Biosciences, Malborough, MA). The 3-Plex Negative Control Probe (dapB, catalog #320871, Advanced Cell Diagnostics) and the positive control probe (polymerase II subunit RPB1, Peptidyl-prolyl cis-trans isomerase B, and ubiquitin C, catalog #300041, Advanced Cell Diagnostics) were processed in parallel with the target probes to confirm assay performance and RNA integrity of the samples.

### Immunohistochemistry

Glp1r-Cre;Ai9 mice were transcardially perfused with chilled 0.9% saline followed by 10% neutral buffered formalin (catalog #SF100-4, Fisher Scientific). Brains were removed and post-fixed 24h at 4°C in 30% sucrose in 10% buffered formalin. Serial coronal sections (40 µM) from frozen brains were taken throughout the GC on a Leica microtome and stored in phosphate buffered saline in 0.3% sodium azide. Sections were rinsed 3X (5 min each) in tris-buffered saline (TBS) and blocked in 5% normal goat serum (catalog #G9023, Millipore) for 1h at room temperature. Next, sections were incubated in rabbit anti-NeuN, (1:1000, catalog #ab177487, Abcam) for 24h at 4°C. Sections were then rinsed 3X for 10 minutes in TBS and incubated in goat anti-rabbit IgG 488 (1:1000, catalog #A11034, Invitrogen) for 2h at room temperature. Finally, sections were rinsed 3X (5 min each) in TBS and 3X in ddH20 prior to mounting sections on slides (catalog #12-550-433, Fisher Scientific). The sections were treated with DAPI-supplemented mounting medium (catalog #OB010020, SouthernBiotech, Birmingham, AL) and cover-slipped.

### Imaging and quantification

Brain regions of interest were identified using the DAPI signal referenced to a mouse brain atlas (Paxinos and Franklin, 2000). Immunofluorescence was detected and imaged using an upright epifluorescent microscope (Nikon Ti2e eclipse) at 10, 20, or 40x magnification. For FISH and immunohistochemical analyses, regions of interest (ROI) were drawn around the GC or NAc to allow for restricted quantification of the fluorescent signal within those areas of interest. For quantification, semi-automated recipes were used to produce counts of the signal(s) of interest based upon fluorescence intensity, size, and localization to a given ROI. Gain, exposure, and light brightness were held consistent across brain regions and animals to allow for accurate comparisons.

### Brain slice electrophysiology

*Ex vivo* brain slice recordings were performed in Glp1r-Cre;Ai9 mice wherein tdTomato expression is directed within cells expressing Glp1r. Mice (n = 11) were anesthetized with isoflurane (4%) and rapidly decapitated. Brains were removed, blocked, and mounted in a vibratome chamber (Leica VT 1200S, Deer Park, IL). Coronal slices (230 µm) containing the GC (coordinates: AP +1.7mm AP, ±3.6mm ML, -3.25mm DV, identified based on anatomical coordinates (Paxinos and Franklin, 2000)) were collected in room temperature artificial cerebrospinal fluid (aCSF) that contained the following (in mM): 126 NaCl, 2.5 KCl, 1.2 MgCl2, 2.4 CaCl2, 1.2 NaH2PO4, 11 d-glucose and 21.4 NaHCO3 (equilibrated with 95% O2–5% CO2). Slices were stored at 32°C in glass vials with carbogen equilibrated aCSF. MK801 (10µM) was added to the cutting solution and for the initial incubation of slices in storage (at least 30min) to block NMDA receptor-mediated excitotoxicity. Following incubation, the slices were transferred to a recording chamber that was perfused with equilibrated aCSF warmed to 34°C (Warner Instruments) at a flow rate of 1.5-3 mL/min. Cells were visualized using an upright microscope (Nikon FN1) equipped with custom built IR-Dodt gradient contrast illumination and a DAGE-MTI IR-1000 camera (Michigan City, IN). Fluorescence was identified using LED epifluorescence illumination with a Texas Red filter cube (excitation: 559nM, emission: 630nM) and detected using a DAGE-MTI IR-1000 camera with sufficient sensitivity in the tdTomato emission range. We targeted and recorded from tdTomato positive (tdTomato+) neurons as GLP-1R+ neurons and nearby tdTomato negative neurons as GLP-1R-null (GLP-1RØ). Whole-cell recordings from layers II/III GC neurons were performed with an Multiclamp 700B amplifier (Molecular Devices, San Jose, CA) in voltage-clamp (Vhold = −60mV) or current clamp mode. Recording pipettes (1.5–2.5MΩ) were filled with internal solution that contained (in mM): 115 potassium methanesulfonate, 20 NaCl, 1.5 MgCl2, 5 HEPES(K), 2 BAPTA, 1–2 Mg-ATP, 0.2 Na-GTP, adjusted to 7.35pH and 275–285mOsM. Liquid junction potential (10mV) was not corrected. Data were low-pass filtered (10kHz) and sampled at 20kHz with pCLAMP 11.1, or at 400Hz with PowerLab (LabChart version 8.1.16, Colorado Springs, CO). To block excitatory and inhibitory fast synaptic transmission, strychnine (1µM), picrotoxin (100µM), and DNQX (10µM) (all from Sigma Aldrich, St. Louis, MO) were added to the aCSF. Current-voltage (I–V) relationships were determined with a series of 10 mV voltage steps (−50 to −140mV) at baseline and during perfusion of the high affinity GLP-1R agonist Exendin-4 (Ex-4; 100nM; catalog #4044219, Bachem, Torrance, CA). The baseline I–V was subtracted from the I–V during Ex-4 to determine the reversal potential of the Ex-4-mediated current. Action potential data were acquired in current-clamp mode by injecting current steps (-50 to +400pA, 500ms). Series resistance was monitored without compensation and remained <20MΩ for inclusion. All drugs were applied by bath perfusion at the indicated concentrations.

### Surgical Procedures

In all surgeries, mice were anesthetized with isoflurane (4% induction, 2% maintenance; IsoFlo®, Patterson Veterinary) in 1 L/min O_2_ and head-fixed in a stereotaxic frame (Model #1900, Kopf, Tujunga, CA) with the torso laying on a water-bath heating pad (38°C). Following injection of local anesthetic (bupivicaine, 5mg/kg, Patterson Veterinary) into the scalp and a midline incision along the skull cap, craniotomies were made bilaterally above the GC (coordinates: +1.7 mm AP, +/-3.4 mm ML)(Paxinos and Franklin, 2000).

For DREADD-based chemogenetic manipulation (Armbruster et al., 2007) of GC GLP-1R+ neurons, AAV8-hSyn-DIO-HA-hM3D(Gq)-IRES-mCitrine (AAV-DIO-hM3Di-mCitrine, 2.3×10^13^vg/ml, 200 nl/hemisphere, catalog #50454-AAV8, Addgene, Watertown, MA) was delivered bilaterally into the GC of Glp1r-Cre mice at the coordinates above at a depth of -3.25mm DV. Following infusion, the micropipette was slowly withdrawn from the brain. The craniotomies were sealed with wax and the scalp closed with Vetbond (3M, Mapelwood, MN).

For pharmacological manipulation of GC GLP-1Rs, bilateral craniotomies were made above the GC and miniature guide cannulae (catalog #C315GMN/SPC, extending +2.25 mm beyond pedestal, P1 Technologies, Roanoke, VA) were implanted at the at the coordinates above at a depth of -2.25 mm DV. Cannulae were secured to the skull by Vetbond followed by dental cement, and dust caps with a 2.25 mm projection wire were inserted.

For both AAV injections and cannulations, mice recovered on a heating pad until ambulatory and then were single-housed. Cannulated mice and AAV-injected mice were given 7 and 14 days, respectively, after surgery to recover prior to the onset of acclimation to behavioral procedures. Meloxicam analgesic (5mg/kg, sc) was administered for at least 3 days following surgery.

### Food intake experiments

Mice were habituated to handling and dark cycle food intake measurements on at least two occasions prior to testing to mimize the effects of stress on food intake. When mice exhibited less than 15% variation in food intake across days, they were considered habituated. Food was removed from cages 2h prior to dark cycle onset, and initial food weight and body weight were recorded at this time. To chemogenetically activate GC GLP-1R+ neurons, Glp1r-Cre animals received ip injections of either sterile saline (vehicle) or the DREADD agonist, JHU37160 dihydrochloride (Bonaventura et al., 2019) (J60, 0.1 mg/kg, catalog #HB6261, HelloBio, Princeton, NJ) 15min prior to dark onset in a counter-balanced fashion. To identify potential off-target effects of J60 on feeding behavior, the same behavioral experiment was conducted in a different subset of Glp1r-Cre mice that did not express AAV-DIO-hM3Di-mCitrine (*i*.*e*., no-DREADDs control).

At conclusion of the DIO behavioral experiments, during perfusion for brain collection (see above), abdominal adipoise tissue was collected and weighed to confirm lean or DIO status. These data are reported as visceral fat over bodyweight.

To pharmacologically activate GC GLP-Rs, mice received intra-GC infusions 30 min-to-1h prior to dark cycle onset. Internal cannulae (extending +3.25 mm beyond pedestal, catalog #C315IMN, P1 Technologies) bilaterally delivered sterile saline (catalog #07-892-4348, Patterson Veterinary, Devens, MA) or Exenatide acetate (Ex-4; catalog #4044219, Bachem) in 0.25µl volume. Pre-weighed food was returned to cages immediately prior to dark cycle onset, and measured at 1, 2, and 22h post-dark onset. To determine the impact of diet-induced obesity (DIO) on GC GLP-1R function, mice were maintained on a low-fat diet (LF, 10% kcal from fat; catalog #D12450Ji, Research Diets, New Brunswick, NJ) or a high-fat diet (HF, 60% kcal from fat; catalog #D12492i, Research Diets). Diets were provided *ad libitum* and mice in each group were matched for age, weight, and sex. Mice were maintained on their respective diets for 6 weeks prior to cannulation and onset of behavioral testing. Behavioral testing followed the same timeline described above.

### Histological verification of cannulae placement and injection accuracy

For histological confirmation of AAV-driven transgene expression and cannula placement, mice were anesthetized and perfused as described above. Brains were removed and post-fixed for 24h at 4°C in 30% sucrose in 10% buffered formalin. Coronal sections (40 µM) were collected through the GC. Sections were slide-mounted with a DAPI-supplemented mounting medium (DAPI Fluoromount-G, catalog #OB010020, SouthernBiotech, Birmingham, AL). Sections displaying AAV-driven transgene expression or cannula tracts were imaged using an upright epifluorescent microscope (Eclipse, Ti2e, Nikon). All mice receiving AAV injections were verified to have region-specific expression in the GC. 6 mice with indwelling cannulae implants were excluded from data analysis due to the cannulae being off-target. This left a total of 45 cannulated c57bl/6j mice for analyses.

### Statistical Analysis

We used a between-subjects design for fluorescent *in situ* hybridization, immunohistochemical, electrophysiological, and behavioral experiments examining the effects of maintainence diet (LF or HF diet) on body weight change. For all other behavioral experiments, within-subjects design was employed. Statistical tests included unpaired two-tailed *t* test, paired two-tailed *t* test, paired one-tailed *t* test, one-, two-, or three way repeated measures analysis of variance (ANOVA), with Šidák or Dunnett’s multiple comparisons *post-hoc* analyses where appropriate as described in the text. Electrophysiological data were tested for normality with the Kolmogorov–Smirnov test. To examine food intake in both male and female mice, the results are presented as chow intake (kcal) divided by body weight (grams) to yield a reflection of food intake corrected by size of the animal. Data are expressed as mean + standard error of the mean (SEM). Differences in means were considered significant when p < 0.05. Statistical tests were performed using GraphPad version 9.2 (GraphPad Software Inc., La Jolla CA, USA).

## Results

### GLP-1Rs are expressed on neurons within the GC

We used RNAscope® to detect *Glp1r* mRNA expression within the GC in C57BL/6J mice. *Glp1r* mRNA was scattered throughout the GC (**Fig 1A,B,C**). We quantified the percentage of DAPI-expressing cells that also expressed *Glp1r* mRNA to determine expression levels which uncovered that roughly 6% of GC cells express *Glp1r* (**Fig 1D,E**). To put this level of GC *Glp1r* mRNA expression in context with other brain-regions known to express GLP-1Rs, we quantified *Glp1r* mRNA from the nucleus accumbens (NAc) in the same brain sections as those sampled from the GC. NAc neurons are known to express GLP-1Rs and antagonism of GLP-1Rs in the NAc influences food intake (Dossat et al., 2011; Alhadeff et al., 2012). *Glp1r* mRNA expression in the GC was comprable to that in the NAc (t(8)=2.179, p=0.061, **Fig 1E;** *n*=4-5 mice, 2-12 sections averaged/mouse).

**Figure 1.**
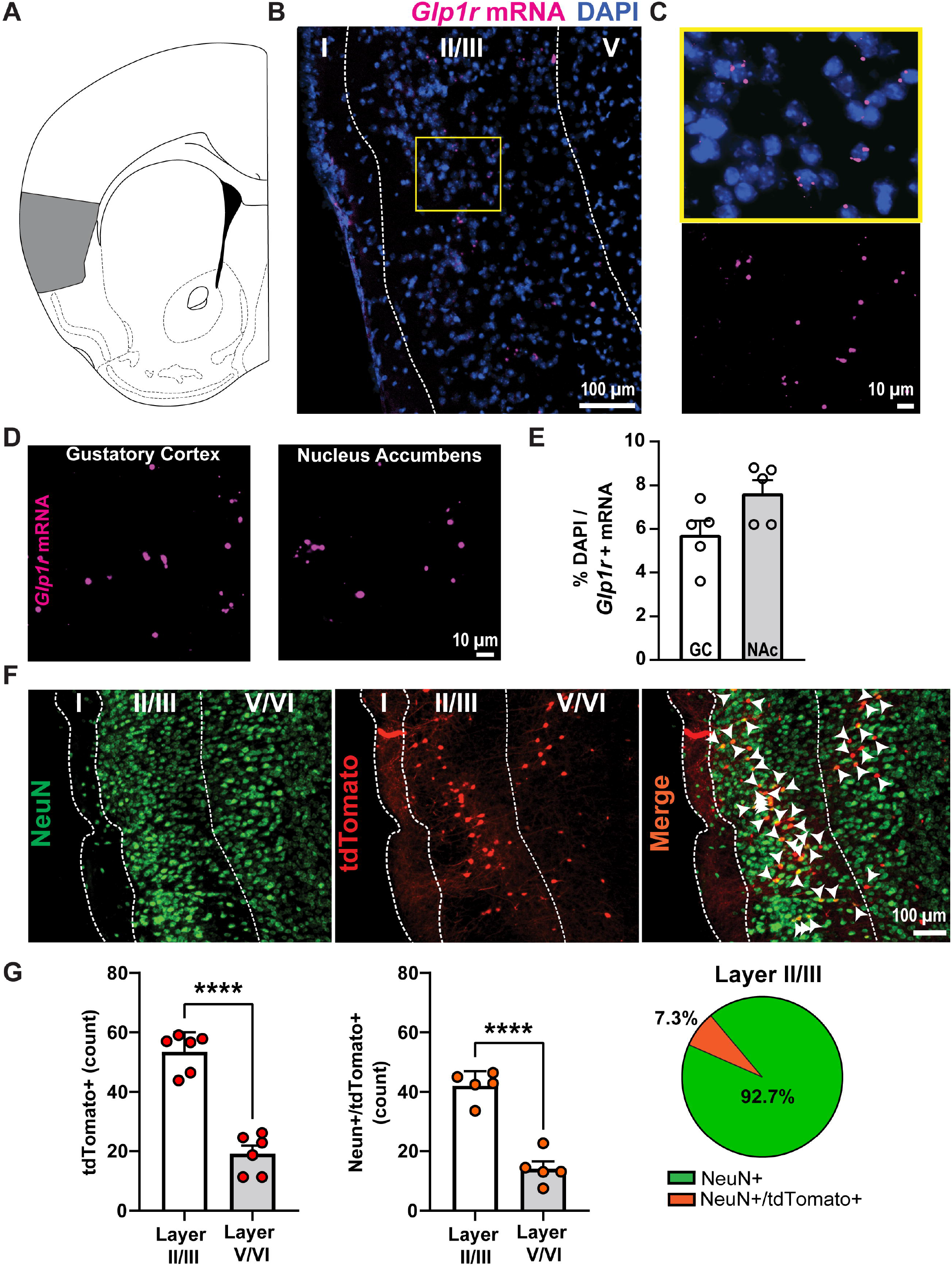
GLP-1Rs are neuronally expressed within the GC. **A**. Localization of the GC (shaded gray) in a coronal section of the mouse brain. **B**. Representative image of G*lp1r* mRNA and DAPI in GC layer II/III. **C**. View of yellow-indicated region in **B** with (top) and without (bottom) DAPI to aid in visualization of G*lp1r* expression. **D**. View of G*lp1r* mRNA in the GC and in the nucleus accumbens (NAc) from the same coronal section and **E**. quantification of *Glp1r* mRNA for each region. n=5 mice, 2-12 sections per mouse. **F**. Representative image of the GC from a Glp1r-Cre;Ai9 mice immunolabeled for the neuronal marker, NeuN, and a merged image showing tdTomato+ somatic colocalization with NeuN (white arrow heads). **G**. Quantification of tdTomato+ neurons, NeuN+/tdTomato+ neurons, and a pie chart depicting total NeuN expression within layer II/III and percent of NeuN+/tdTomato+ cells. Data are mean +SEM. n=6 mice, 5-10 sections per mouse. ****p<0.0001 layer II/III vs. V/VI.

Next, we paired immunohistochemistry and genetically-driven fluorophores in Glp1-Cre mice crossed with the Ai9 tdTomato reporter line (Glp1r-Cre;Ai9) to determine if there is layer specificity of GC GLP-1Rs and, seperately, if they are neuronally-localized. Sections from Glp1r-Cre;Ai9 mice revealed significantly higher expression of GLP-1R+ neurons (tdTomato+) within GC layers II/III vs. layers V/VI (t(10)=9.072, p<0.0001, **Fig 1F,G**), and higher expression of NeuN+ and GLP-1R+ expression in layers II/III vs. layers V/VI (t(8)=8.507, p<0.0001). Of the NeuN+ signal within layers II/III, we found that 7.3% of the NeuN+ signal also expressed GLP-1R (tdTomato+) – a number comprable to that revealed in our prior RNAscope experiment (**Fig 1E**). Together, these data indicate the GC GLP-1Rs are localized to neurons and that GLP-1R neurons are most abundant in the dense cell layers II and III of the GC.

### GC GLP-1R+ neurons are depolarized by the selective GLP1-R agonist, Exendin-4

To determine whether activation of GLP-1Rs functionally modulates GC neurons, we performed whole-cell voltage-clamp recordings from layer II/III GC GLP-1R+ and GLP-1RØ neurons visualized in brain slices from Glp1r-Cre;Ai9 mice (**Fig 2A**). Application of the selective GLP-1R agonist Ex-4 (100 nM) produced an inward current in GLP-1R+ neurons (Ex-4 holding current vs. baseline, paired two-tailed t-test (t(9)=3.165, p = 0.012), but not in neighboring GLP-1RØ neurons (Ex-4 holding current vs. baseline, paired two-tailed t-test (t(8)=1.936, p = 0.089) (**Fig 2B,C**). The Ex-4 mediated currents in GLP-1R+ neurons reversed at -85 ± 3.2 mV, which is near the expected reversal potential for potassium, and was associated with an increase in membrane resistance (baseline = 106 ± 11 MΩ vs. Ex-4 = 121 ± 12 MΩ, paired two-tailed t-test (t(9)=4.302, p = 0.002)), indicating a closure of ion channels.

**Figure 2.**
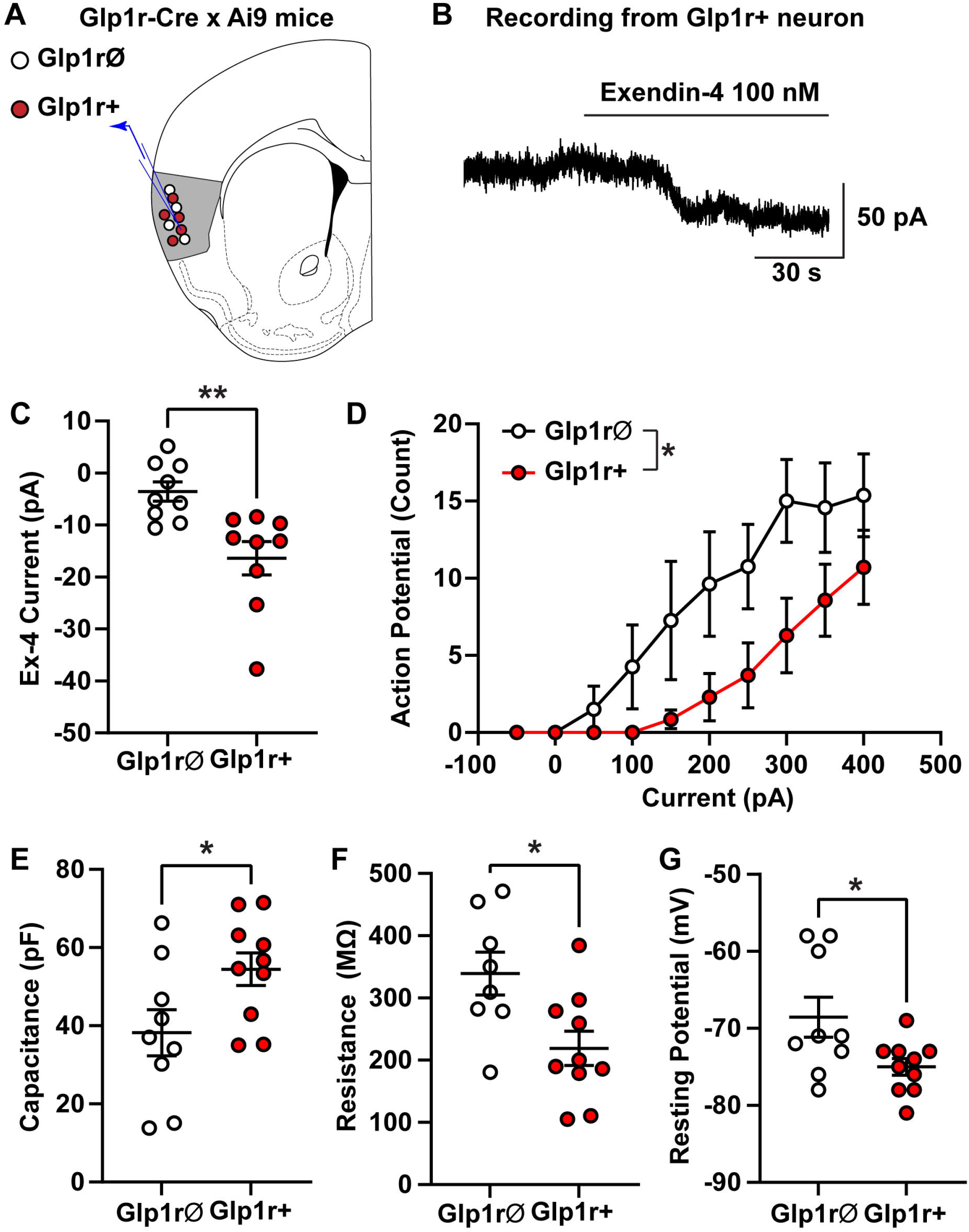
GC GLP1-R+ neurons are depolarized by Ex-4 and exhibit distinct electrophysiological properties vs. neighboring GLP-1RØ neurons. **A**. Whole-cell voltage-clamp recordings were made from GC layer II/III Glp1r+ and Glp1rØ neurons from Glp1r-Cre;Ai9 mice. **B**. Example recording from Glp1r+ neuron showing slow inward current induced by bath application of Ex-4 (100 nM). **C**. Ex-4 mediated inward currents were larger in Glp1r+ neurons compared to Glp1rØ neurons. **D**. Number of action potentials elicited with increasing current steps was different in Glp1r+ and Glp1rØ neurons (two-way ANOVA, F(1,13)=5.95, p=0.0298). Glp1r+ and Glp1rØ neurons had different intrinsic properties, including **E**. capacitance, **F**. membrane resistance and **G**. resting membrane potential. Data are mean ± SEM. n=7-10/group. *p<0.05, **p<0.01 by unpaired t-test.

Additionally, GLP-1R+ and GLP-1RØ neurons exhibited different intrinsic properties. GLP1-R+ neurons had a higher membrane capacitance (unpaired two-tailed, t-test: t(17)=2.284, p=0.036), lower membrane resistance (t(16)=2.778, p=0.013), and more hyperpolarized resting membrane potential (t(17)=2.384, p=0.03) as compared to neighboring GC GLP-1RØ neurons (**Fig 2E,F,G**). Under current clamp conditions, the number of action potentials (APs) produced by depolarizing currents was less in GLP-1R+ neurons compared to GLP-1RØ neurons (two-way ANOVA, F(1,13)=5.95, p=0.029) (**Fig 2D**). These data suggest that GLP-1R+ and GLP-1RØ neurons are functionally different populations of neurons, and that GLP-1R+ neurons are, most importantly, that they are depolarized by GLP-1R agonism.

### GC GLP-1R+ neurons and GC GLP-1Rs influence food intake

Having identified GLP-1Rs in the GC, and having found that these receptors are functional on the neurons (capable of driving depolarization upon their agonism), we next sought to perform two seperate sets of studies to examine the possible causal influence of GC GLP-1Rs on food intake. These included both chemogenetic manipulation of GC GLP-1R+ neurons, and pharmacological interrogation of the GC GLP-1Rs. In both studies, we examined dark-cycle intake of standard chow/food pellets as a simple assay for homoestatic food intake.

In the first experiment, we chemogenetically activated GC GLP-1R+ neurons via administration of the DREADD ligand J60 (0.1 mg/kg, ip.) in Glp1r-Cre mice with bilateral expression of AAV-DIO-hM3Di in the GC (**Fig 3A,B,C**). There was no significant effect of sex (two-way ANOVA: sex: F(1,5)=0.089, p=0.778; DREADD agonism: F(1,5)=3.93, p=0.104), as such data were pooled across sexes. Chemogenetic activation of GC GLP-1R+ neurons significantly reduced chow intake at 1h post-dark onset (one-tailed, paired t-test; t(6)=2.066, p=0.042, **Fig 3B left**), with no effect at 2 or 24h (data not shown). Importantly, a seperate group of Glp1r-Cre mice that did not express AAV revealed that J60 (same dose as above) did not influence chow intake (two-tailed, paired t-test, t(6)=0.416, p=0.692, **Fig 3B right**), indicating the effect of J60 on food intake is specific to DREADD activation.

**Figure 3.**
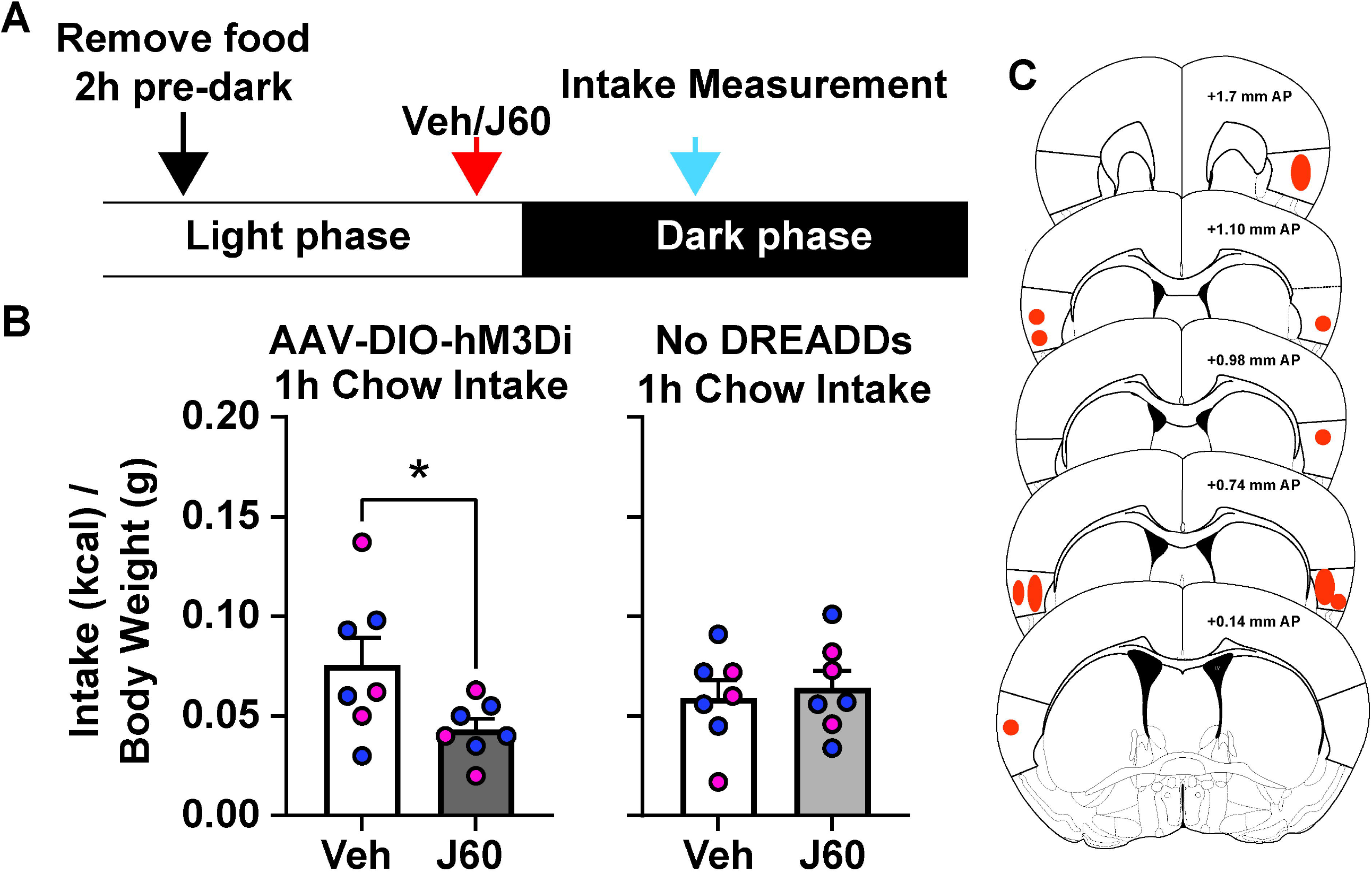
Chemogenetic activation of GC GLP-1R+ neurons reduced dark cycle chow intake. **A**. Schematic of experimental timeline. **B**. Effect of DREADD ligand J60 (0.1mg/kg, ip) on dark cycle chow intake in Glp1r-Cre mice bilaterally expressing GC AAV-DIO-hM3Di-mCitrine, and in Glp1r-Cre mice that do not express DREADD receptors. Mean + SEM of chow consumed (corrected to kcal consumed divided by body weight). n=7/group (3 females, 4 males), blue circles = males, pink circles = female. *p<0.05 vs. vehicle. **C**. Shaded regions illustrating representative spread of AAV-DIO-hM3Di-mCitrine expression within the GC. Anterior-posterior (AP) coordinates are relative to bregma.

In the second experiment, we delivered the GLP-1R agonist, Ex-4, at a range of doses in a counter-balanced order over different days into the GC of C57BL/6J mice via bilateral indwelling cannulae. Based upon our anatomical data (**Fig 1G**), we attempted to target GLP-1Rs within GC layers II/III due to the high numbers of GLP-1R+ neurons in these layers. There was no significant effect of sex (two-way ANOVA, sex: F(1,8)=0.562, p=0.475; Ex-4: F(2.748,21.99)=5.392, p=0.007), and therefore going forward data were pooled across sexes. We found that intra-GC Ex-4 significantly reduced dark cycle chow intake at 2h post-dark onset (one-way ANOVA: F(2.718,24.46)=5.693, p=0.005). Significant reductions in chow intake were uncovered following infusions of both 0.03 and 0.1 ug of Ex-4 (Dunnett’s *post-hoc*: t(9)=3.067, p=0.034, t(9)=3.72, p=0.012, respectively, **Fig 4B**). Taken together, the results of both the chemogenetic and pharmacological experiments support a role for GC GLP-1Rs in influencing homeostatic food intake.

**Figure 4.**
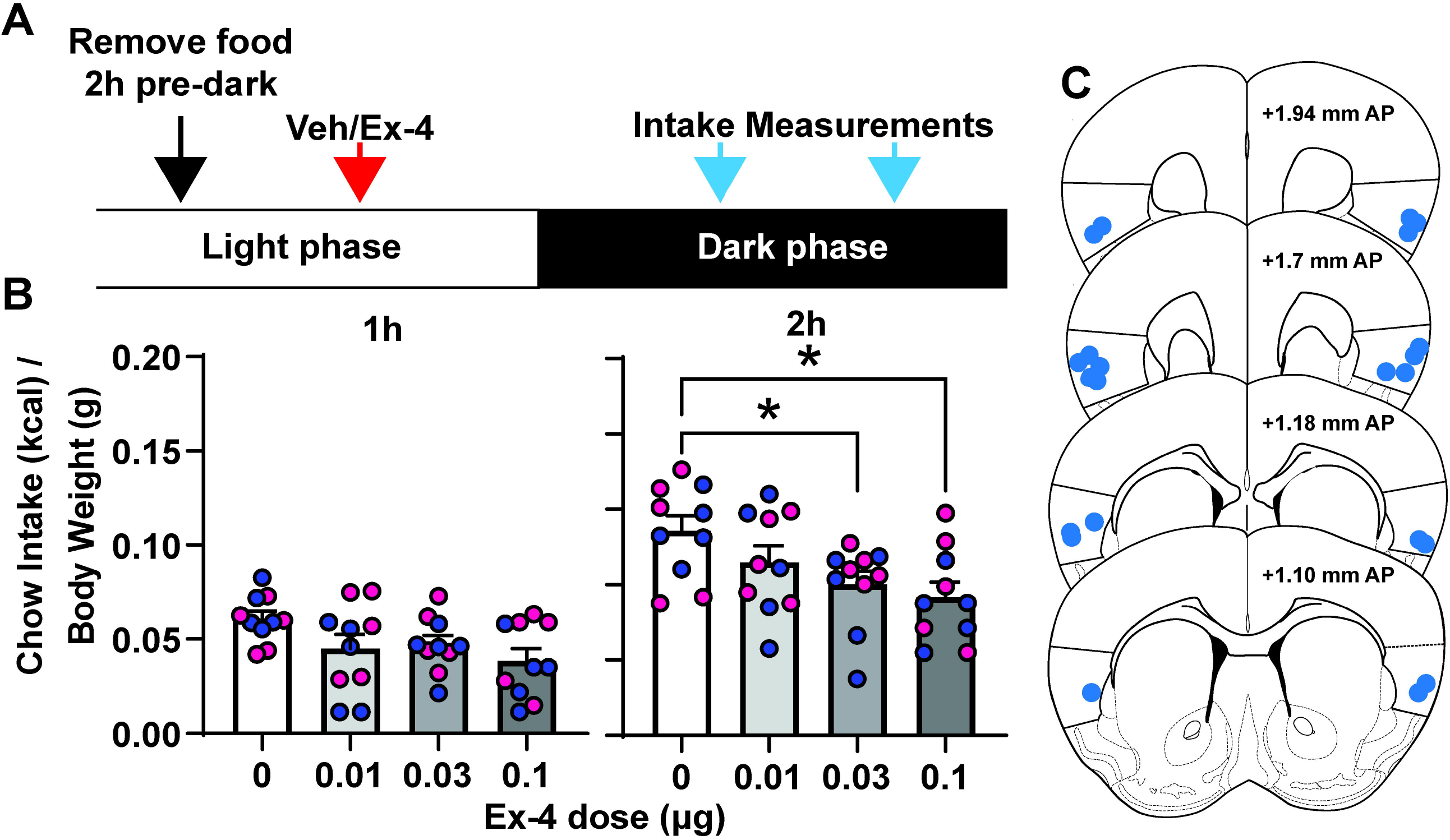
Pharmacological activation of GC GLP-1Rs reduced dark cycle chow intake. **A**. Schematic of experimental timeline. **B**. Effect of intra-GC Ex-4 on chow intake at 1 and 2h into the dark cycle. **B**. Mean + SEM of chow consumed (corrected to kcal consumed divided by body weight). n=7/group (3 females, 4 males), blue circles = males, pink circles = female. *p<0.05 vs. vehicle. **C**. Representative cannula placements of mice. Anterior-posterior (AP) coordinates are relative to bregma.

### Effect of GC GLP-1R agonism on palatable food intake in obese mice

While homeostatic food intake reflects a basic desire to consume food, ingestive behavior is a complex process mediated by a variety of both internal and external factors. Therefore, we next explored whether the above effects of GC GLP-1Rs on food intake extended into influencing the consumption of palatable foods and how this may be also modulated by the metabolic state of the mice. To accomplish this, C57BL/6J mice were given *ad libitum* access to either low fat (LF; 10% kcal fat) or high fat diet (HD; 60% kcal fat) for 6 weeks to induce diet-induced obesity (DIO) or not in the HD or LD groups, respectively. Consistent with prior reports, we observed an increase in body weight in HF-maintained animals soon upon diet onset (e.g., (Williams et al., 2011; Honors et al., 2012; Salinero et al., 2018; Maric et al., 2022)). There was a significant main effect of diet, time on diet, and an interaction between diet x time on change in body weight (two-way ANOVA, diet: F(1,33)=30.06, p<0.0001; time: F(1.751,57.80)=66.73, p<0.0001; interaction: F(5,165)=26.99, p<0.0001; **Fig 5A**). Also as expected (Hwang et al., 2010; Carlin et al., 2016; Salinero et al., 2018; Maric et al., 2022), we observed a robust sex-difference in DIO expression, wherein females exhibited less weight gain as compared to males when maintained on the same HF diet (**Fig 5B**). This was confirmed via fat pad analysis at the conclusion of the experiment (**Fig 5G**). We compared males’ and females’ change in body weight over the 6 weeks of HF consumption, and identified a significant main effect of sex (F(1,20)=7.111, p=0.0148), a significant effect of time on diet (F(1.598, 31.95)=70.24, p<0.0001), and a significant interaction between sex x time on diet (F(5, 100)=4.99, p=0.0004). An examination of the body weight change in HF males and females only at week 6 revealed a significant effect of sex (two-tailed, unpaired t-test, t(20)=4.023, p=0.0007, **Fig 5C**).

**Figure 5.**
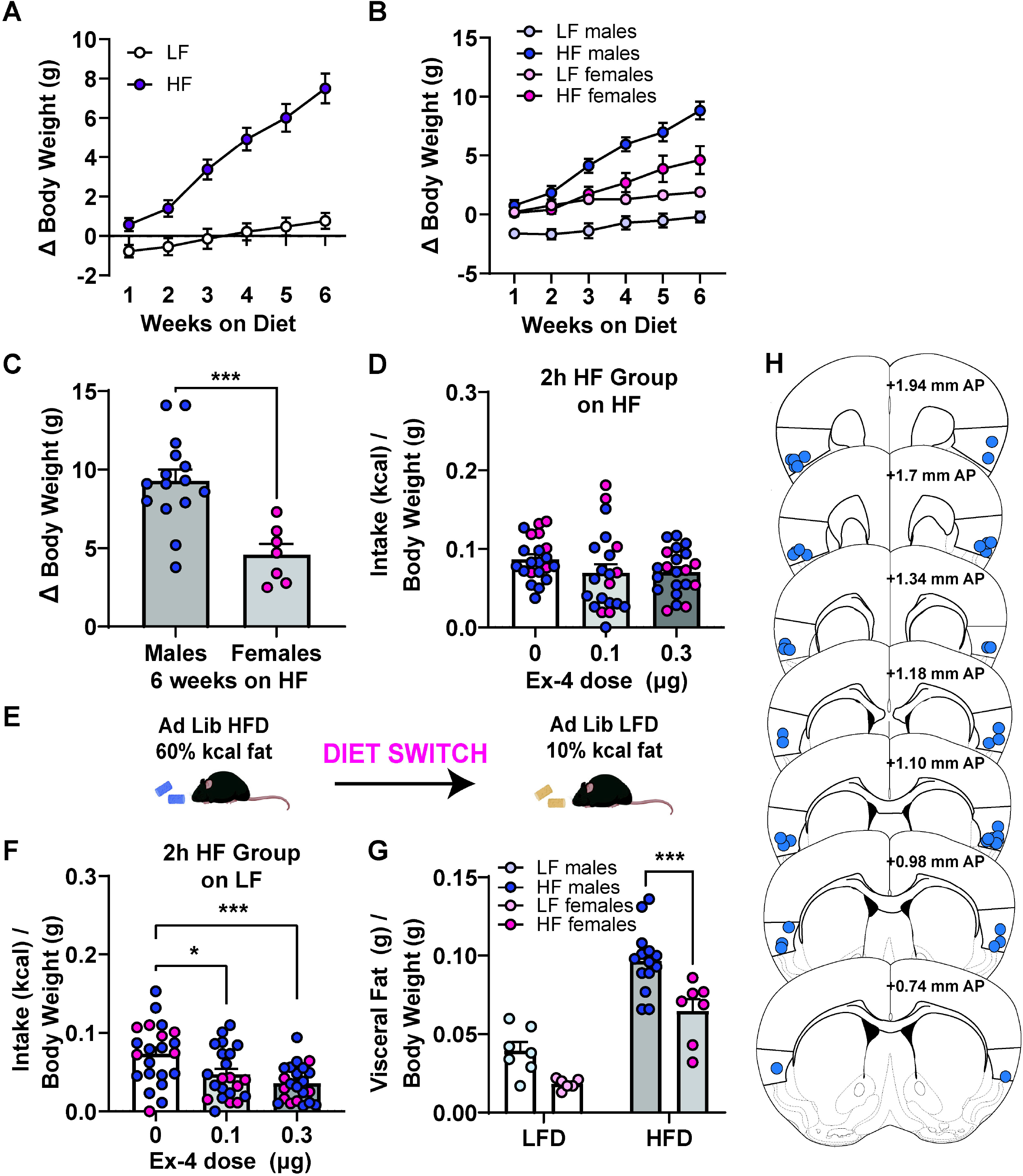
Impact of intra-GC Ex-4 in context of diet induced obesity. **A**. Weekly change in body weight in mice on the LF or HF diet during the development of diet-induced obesity (DIO) over six weeks. **B**. Weekly change in body weight from **A**, divided by sex. **C**. Comparing the average change in body weight on the 6^th^ week of HF maintenance in males and females. ***p<0.001 males vs. females. **D**. Intake (kcal/body weight) of HF by the HF-maintained group, 2h into dark cycle following intra-GC Ex-4. **E**. Schematic of experimental timeline, animals on HF acutely switched to LF. **F**. LF intake kcal/body weight) of the HF-maintained group, 2h into the dark cycle following intra-CG Ex-4. *p<0.05, ***p<0.001 vs. 0 µg Ex-4. **G**. Visceral fat (g) calculated over body weight (g) at the end of the experiment (see Methods). ***p<0.005 HF males vs. HF females. **H**. Representative cannula infusion tip placements of mice included in panels **D**,**F**,**G**. n=13-22 mice/diet condition.

After six weeks on the respective diets and having confirmed a DIO state, the GC was cannulated and mice received intra-GC Ex-4 following the same treatment timeline as above. There was no effect of sex (two-way ANOVA, sex: F(1,20)=1.483, p=0.238), as such data are pooled across males and females. The HF group consuming the HF diet exhibited no Ex-4-induced reductions in food intake at 2h into the dark cycle (F(1.404, 28.08)=2.199, p=0.142; **Fig 5D**). After an acute diet switch (cycle of 3 days on LF followed by 4 days on HF to maintain body weight, **Fig 5E**), we examined intra-GC Ex-4 induced responses in the HF group consuming a LF diet. There was no effect of sex (two-way ANOVA, sex: F(1,20)=0.200, p=0.659), as such data are pooled across males and females. In this case, there was a significant main effect of Ex-4 to reduce LF diet intake at 2h into the dark cycle (F(1.631,34.26)=9.801, p=0.001; **Fig 5F**). *Post-hoc* tests revealed significant effects of both 0.1 and 0.3 µg Ex-4 (t(21)=2.519, p=0.037; t(21)=4.307, p=0.001; respectively). *Post-mortem* analyses confirmed the mice contributing to tehse data had cannulae localized within the GC (**Fig 5H**). Taken together with the influence of GC GLP-1R modulation of intake of standard lab chow (**Fig 4**), these results suggest that GC GLP-1Rs may influence both homeostatic (eating standard lab chow) and hedonic (preferring HF or LF) food intake. Further, metabolic state, in this case obesity, influences the consequences of GC GLP-1R signaling on subsequent food intake.

## Discussion

The GC integrates taste, valence, and interoceptive information. Cells within the GC are well-known to exhibit tastant-evoked responses relating to tastant identity as well as the palatability of tastants (Katz et al., 2001; Small et al., 2003; Stapleton et al., 2006; Jezzini et al., 2013; Maier and Katz, 2013; Fletcher et al., 2017; Chen et al., 2021). Yet, it is interesting to note that some GC cells do not respond to tastants and there are both layer and cell type-specific responses within the GC (Dikecligil et al., 2020). This heterogeneity within the GC underscores the need for understanding the role of distinct neuronal populations within this brain area that may contribute to behavior, especially ingestive behaviors. Indeed, in addition to taste-processing, GC activity is modulated by prandial state wherein activity is highest when hungry and reduced when satiated (Small et al., 2001; Saper, 2002; Hinton et al., 2004; de Araujo et al., 2006; Van Bloemendaal et al., 2014). Additionally, preclincial work demonstrated that GC neurons are sensitive to metabolic state (*i*.*e*., fasted or fed) and respond to food-predicting cues more robustly when fasted vs. fed (Livneh et al., 2017, 2020). In the present study, we demonstrate the presence and functionality of a previously unknown population of GLP-1Rs within the GC.

### Localization of functional GLP-1Rs in the Gustatory Cortex (GC)

To our knowledge, we provide the first evidence for GLP-1R expression within the GC. We found that approximately 5% GC cells express *Glp1r* mRNA – a finding corroborated when crossing Glp1r-Cre mice with Ai9 Tdtomato reporter mice. While this is a relatively small proportion of cells, we found this expression level was comparable to that in the NAc, a region in which GLP-1R agonism robustly reduces food intake (Dossat et al., 2011, 2013; Alhadeff et al., 2012). Prior work in a different brain regions uncovered evidence for expression of GLP-1Rs on astrocytes (Reiner et al., 2016). Importantly, through immunohistochemistry with the neuronal marker NeuN in Glp1r-Cre;Ai9 mice, we verified that GLP-1R+ cells in the GC appear to be neurons, consistent with the majority of research in other brain regions (Tauchi et al., 2008; Richard et al., 2014; Fortin et al., 2020; Zeng et al., 2021).

The differing GC layers subserve distinct roles in sensory processing. For instance, layers V/VI contain a large number of taste-sensitive neurons (Kayyal et al., 2019; Dikecligil et al., 2020). The bulk of GLP-1R+ neurons were found within layers II/III of the GC, with GLP-1R+ neurons detected within layers V/VI but a significantly lower level than layers II/III. While the synaptic organization of the GC is a topic of much interest (e.g., (Haley et al., 2016, 2020)), it has received considerably less attention that the synaptic organization of many other cortices. Future research will be needed to define the synaptic/circuit contributions of GLP-1R signaling on GC neurons, especially concerning if and how signaling via layers II/III GC neurons may possibly influence taste processing among the more deep taste-responsive neurons found in layers V/VI.

Perhaps informing the above goal to some extent, we found that application of Ex-4 produced a slow inward current in GLP-1R+ neurons, which was similar in amplitude to inward currents produced by GLP-1 in other areas of the brain (Cork et al., 2015). However, GLP-1Rs may depolarize GC neurons using different intracellular mechanisms and ion channels, since currents in GC neurons reversed near the potassium equilibrium potential and were associated with a decrease in conductance (indicating closing of potassium channels), wheras GLP-1R mediated currents in the PVN, BNST and hippocampus were associated with an increase in conductance. In addition, GLP-1R+ neurons had different intrinsic properties compared to neighboring GLP-1RØ neurons, suggesting that while GLP-1R+ neurons are localized to the densely-packed layer II/III of the GC, they represent a unique population of neurons within the GC.

### A role for GC GLP-1Rs in ingestive behavior

To examine the possible functional role of GC GLP-1R+ neurons in food take, and seperately, GLP-1R signaling within GC neurons, we used both chemogenetics and pharmacology in separate groups of mice. First, we used chemogenetic targeting of GLP-1R+ neurons using Glp1r-Cre mice expressing excitatory DREADD in their GC. When GC GLP-1R+ neurons were activated just prior to the onset of the dark cycle, chow intake was significantly reduced as compared to the control condition. This indicates that this cell population influences ingestive behavior. These results are in agreement with recent work from several groups supporting roles for subsets, sometimes sparse subsets, of GC neurons (*e*.*g*., PKC-δ+, Nos1+,CaMKIIα+) to influence ingestive behavior (Stern et al., 2020, 2021; Wu et al., 2020; Zhang-Molina et al., 2020).

We followed up our chemogenetic findings by exploring how pharmacological manipulation of *specifically* the receptors themselves would influence homeostatic feeding. We found that activation of GC GLP-1Rs with Ex-4 significantly reduced dark cycle chow intake. This was achieved within a dose-range that has been reported for other brain regions (Swick et al., 2015; Ong et al., 2017; López-Ferreras et al., 2018; Reiner et al., 2018). These data indicate that GLP-1R signaling on GC neurons contributes to the control of homeostatic food intake, consistent with work in other brain areas (Tang-Christensen et al., 1996; Dossat et al., 2011; Kanoski et al., 2011; Hsu et al., 2015; Terrill et al., 2019). We observed similar outcomes in males and females to intra-GC GLP-1R activation, which is consistent with prior research on the lateral ventricle (Richard et al., 2016), but inconsistent with effects within the lateral hypothalamus that did uncover sex differences (López-Ferreras et al., 2019). These distinct effects are likely mediated by the brain region targeted, since activation of GLP-1Rs can yield differing effects depending on brian-region (reviewed in (Williams, 2022)).

Pathophysiological states impact GC activity. Specifically, the GC of obese individuals displays greater activation in response to taste stimuli and food-associated cues than non-obese individuals (Van Bloemendaal et al., 2014; Avery et al., 2017; Bohon, 2017; Li et al., 2017). The GLP-1 system itself is also impacted by obesity. Maintenance on high fat diet reduces GLP-1R expression in a brain region-specific manner (Mul et al., 2013) and impairs anorexigenic effects of peripherally administered GLP-1R agonists (Williams et al., 2011; Duca et al., 2013; Mul et al., 2013; Mella et al., 2017). In the present study, we aimed to determine the effect of diet-induced obesity (DIO) on the effects of GC GLP-1Rs on food intake. Male and female mice were maintained on a control LF diet or a HF diet for 6 weeks to induce the DIO phenotype. We found that male and female mice maintained on a HF diet were insensitive to the effects of GC GLP-1R activation, with no sex differences observed within this group. However, following an acute switch from a HF to a LF diet (4 days on HF, 3 days on LF), DIO mice exhibited significant reductions in LF intake following GC GLP-1R activation. Taken together, these data suggest that GC GLP-1Rs remain functional in DIO conditions, yet palatability may impact the ability of GC GLP-1R agonism to reduce intake. It is important to note that we did not examine expression levels of GLP-1Rs in the DIO condition, nor did we directly examine the sensitivity of these receptors. Future investigations examining the impact of DIO on GLP-1R sensitivity and expression in the GC, and its functionality, will be important.

### Conclusion

The present study demonstrates that a population of GC neurons express GLP-1Rs, these neurons are depolarized by agonism of GLP-1R, and these receptors can influence food intake upon their activation. This work provides a foundation for future work to resolve the upstream (where does GLP-1 arise from to reach the GC?) and downstream (where do GC GLP-1R neurons innervate to influence behavior?) circuitry for the GC GLP-1R system. It would be of interest to investigate the possible role, or lack thereof, for GC GLP-1Rs in the known effects of GLP-1 mimetics in managing obesity (Wilding et al., 2021). Further, given the extensive connectivity of the insular cortex, it is important to note that the GC GLP-1R system may have implications beyond ingestive behavior and metabolic conditions. Indeed, awareness for a potential modulatory role of Glp-1 mimetics in a variety of human conditions is increasing (e.g., (Brauer et al., 2020)).

## Acknowledgements

We thank doctors A. Spector, A. Fontanini, and S. Munger for helpful discussions during the course of the project. This work was supported by University of Florida Research Opportunity Seed Fund 00130743 to A.M.D and D.W.W., R01DA047978 to E.S.L., and R01DC014443, R01DA049545, and R01DC016519 to D.W.W.

